## Supplemental Table and Fig for "MDC1 mediates Pellino recruitment to sites of DNA double-strand breaks"

### Supplementary Table S1 – ITC binding data

| *Pellino 1 FHA* | | | | | |
|---|---|---|---|---|---|
| **Peptide** | **Kd (uM)** | **N** | **ΔH (kJ/mol)** | **ΔG (kJ/mol** | **TΔS (kJ/mol K)** |
| MDC1-pT699 | 2.01 | 0.36 | -77.5 | -32.5 | -44.9 |
| MDC1-pT719 | 1.14 | 0.78 | -91.2 | -34.0 | -57.3 |
| MDC1-pT752 | 1.14 | 0.74 | -77.2 | -33.9 | -43.3 |
| MDC1-pT765 | 0.23 | 0.62 | -92.1 | -38.0 | -54.1 |
| IRAK1-pT141+3Y | 0.42 | 0.70 | -72.6 | -36.5 | -36.1 |
| H2AX-pS139 | N.D.B. | - | - | - | - |
| *Pellino 2 FHA* | | | | | |
| **Peptide** | **Kd (uM)** | **N** | **ΔH (kJ/mol)** | **ΔG (kJ/mol** | **TΔS (kJ/mol K)** |
| MDC1-pT699 | 1.12 | 0.55 | -83.4 | -34.0 | -49.4 |
| MDC1-pT719 | 1.22 | 0.52 | -145 | -33.8 | -111 |
| MDC1-pT752 | 1.06 | 0.69 | -118 | -34.1 | -83.7 |
| MDC1-pT765 | 0.37 | 0.69 | -137 | -36.7 | -100 |
| IRAK1-pT141+3Y | 0.70 | 0.59 | -108 | -35.2 | -73 |
| H2AX-pS139 | N.D.B. | - | - | - | - |
| *MDC1 BRCT-r* | | | | | |
| **Peptide** | **Kd (uM)** | **N** | **ΔH (kJ/mol)** | **ΔG (kJ/mol** | **TΔS (kJ/mol K)** |
| H2AX-pS139 | 0.85 | 0.60 | -58.2 | -34.6 | -23.5 |

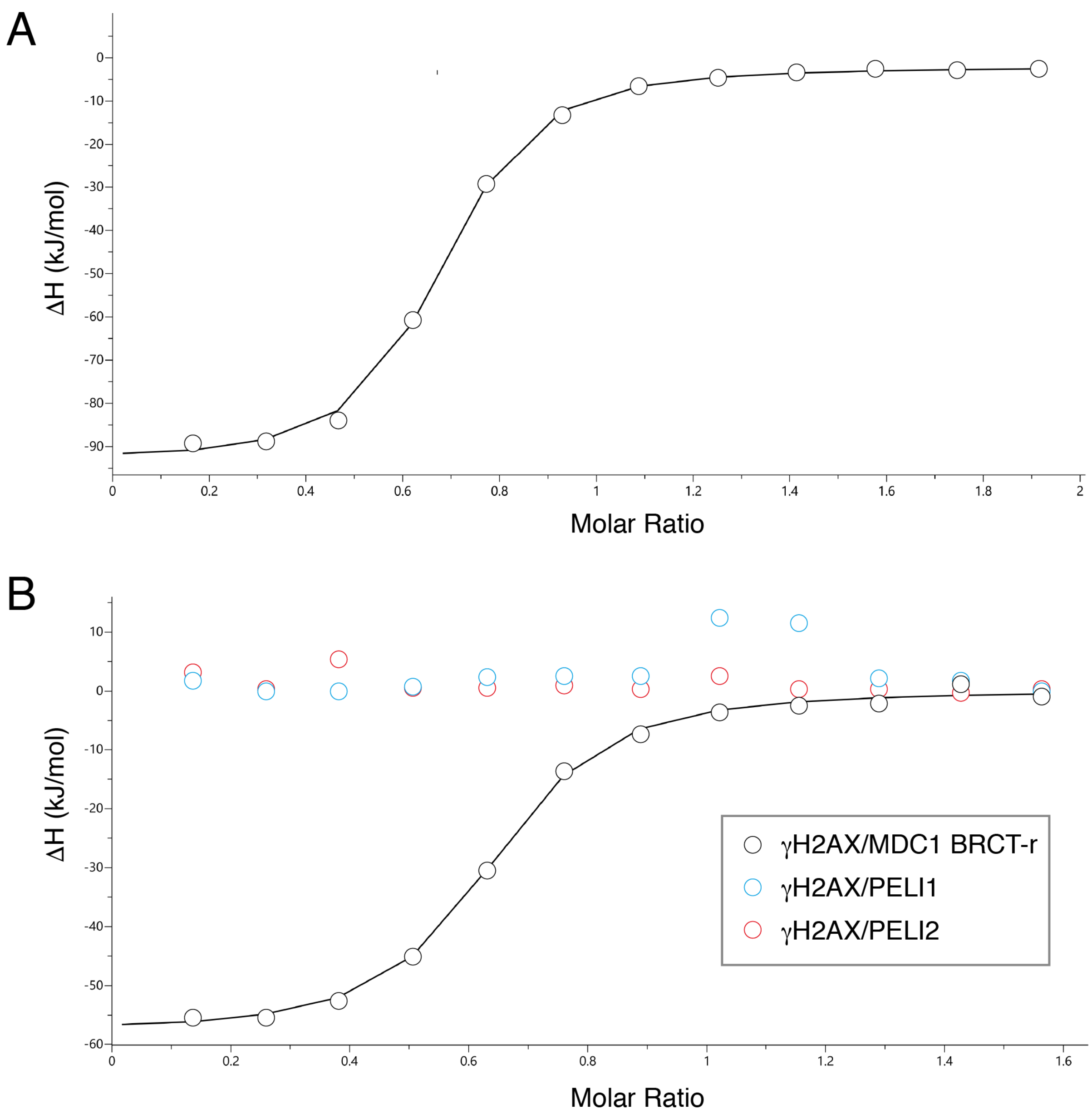

***Supplementary Figure S1 – Phospho-dependent Pellino binding A.*** *ITC titration of PELI1 (13-273) with MDC1-pT765 (Site 4) phosphopeptide* ***B****. ITC titrations of PELI1 (13-273), PELI2 (15-273) and MDC1 BRCT-r (Stucki et al., 2005) with γH2AX phosphopeptide shows tight binding by the MDC1 fragment but no detectable interactions with either PELI1 or 2 FHA regions.*

**Supplementary Table S2 – Crystallographic statistics**

| **Data collection/processing** | |
|---|---|
| Beamline | Diamond Light Source IO2 |
| Wavelength (Å) | 0.9795 |
| Space group | $P2_12_12_1$ |
| Cell parameters (Å) | 52.9, 75.1, 165.1 |
| Cell parameters (°) | 90.00, 90, 90.00 |
| Resolution range (Å) | 68.35 – 2.74 (2.78 – 2.74)* |
| Number of observations | 224435 (6165) |
| Unique reflections | 17750 (727) |
| Completeness (%) | 98.3 (80.7) |
| Mean I/I(σ) | 8.0 (0.9) |
| Multiplicity | 12.6 (8.5) |
| $R_{merge}$ | 0.236 (2.387) |
| $R_{meas}$ | 0.264 (2.449) |
| $R_{pim}$ | 0.069 (0.873) |
| $CC_{1/2}$ | 0.997 (0.407) |
| **Refinement** | |
| Protein atoms | 3732 |
| Waters | 78 |
| Sulphate ions | 5 |
| $R_{cryst}$ | 0.25 (0.41) |
| $R_{free}$ | 0.27 (0.45) |
| RMSD bond-lengths (Å) | 0.013 |
| RMSD bond angles (°) | 1.8 |
| *Ramachandran* | |
| Most Favoured % | 96.9 |
| Additional allowed % | 3.1 |
| Disallowed % | 0 |

* Highest resolution shell in parenthesis.
